# Consensus outlier detection in survival analysis using the rank product test

**DOI:** 10.1101/421917

**Authors:** Eunice Carrasquinha, André Veríssimo, Susana Vinga

## Abstract

Survival analysis is a well known technique in the medical field. The identification of individuals whose survival time is too short or to long given their profile, assumes great importance for the detection of new prognostic factors. The study of these outlying observations have gained increasing relevancy with the availability of high-throughput molecular and clinical data for large cohorts of patients. Several methods for outlier detection in survival data have been proposed, which include the analysis of the residuals, the measurement of the concordance c-index, and methods based on quantile regression for censored data. However, different results are obtained depending on the type of method used. In order to solve the disparity of results we proposed to apply the Rank Product test. A simulated dataset, and two clinical datasets were used to illustrate our proposed consensus outlier detection method, one from myeloma disease and the other from The Cancer Genome Atlas (TCGA) ovarian cancer. Finally, the Rank Product with multiple testing corrections was performed in order to identify which observations have the highest rank amongst the methods considered. Our results illustrate the potential of this consensus approach for the automated retrieval of outliers and also the identification of biomarkers associated with survival in large datasets.

## 1 Introduction

Survival analysis is a statistical technique widely used in many fields of science, in particular in the medical area, and which studies the time until an event of interest occurs. The event may be death, the relapse of a tumour, or the development of a disease. The response variable is the time until that event, called survival or event time, which can be censored, i.e. not observed on all individuals present in the study.

There are different ways of modelling this type of data, one of most widely used due to its flexibility is the Cox proportional hazards regression model [1]. This is a semi-parametric model because the baseline hazard does not need to be specified. One of the problems of this technique is the fact that a single abnormal observation can affect the parameter estimates. To overcome this issue it is important to study mechanisms to identify individuals who lived too long or too short, given their covariates.

The detection of outliers in survival data has gained great importance due to the fact that the identification of individuals with survival time too high or too short can lead in the medical field to the detection of new prognostic factors [2]. The first attempt to analyse and identify outliers was based on residuals. In this context, graphical methods based on the analysis of martingale, score and deviance residuals were proposed [3]. Nardi and colleagues [2] proposed two new types of residuals: the log-odds and normal deviate residuals, with theoretical properties and empirical performance appealing for outlier detection.

More recently, three outlier detection algorithms for censored data were presented [4]: the residual-based, boxplot, and scoring algorithms, all based on quantile regression, which is robust to outliers [5]. Alternative methods for outlier detection based on the concordance c-index were also proposed [6]: the one-step deletion (OSD) and bootstrap hypothesis testing (BHT) strategy, subsequently improved with a dual bootstrap hypothesis testing (DBHT) version [7].

The aim or this paper is to review some of the methods usually applied for outlier detection in survival data and present an ensemble approach that can combine the results obtained by each method, since each usually provides distinct and sometimes contradictory results. In fact, to overcome these differences, we proposed to use a non-parametric method, the Rank Product (RP) test, to identify the outliers that are consistently highly ranked in the outlier detection methods described. In this context, the RP test seems a promising approach to identify outliers in a model-based context. In particular model-based outlier detection methods provide a structured framework to identify abnormal cases, i.e., those who significantly deviate from what would be expected.

All the R scripts and datasets are available at http://web.ist.utl.pt/susanavinga/outlierRP for sake of reproducibility. This includes R Markdown (RMD) files that can be run to replicate all the obtained results.

The outline of this paper is as follows. In Section 2, Survival Regression models, three different approaches to model survival data are presented. In Section 3, several outlier detection methods and the RP test are presented. In Section 4 application examples with simulated data and two clinical datasets are presented. Finally, conclusions and future work are addressed to Section 5.

## 2 Survival Regression models

In this section three different approaches to model survival data are presented. First is introduced the Cox regression model proposed in [1] and then two alternative proposals, robust version on the Cox regression model ([10]) and censored quantile regression ([16]), are presented.

### 2.1 Cox regression model

The Cox regression model is one of the most used methods in Survival analysis [1]. It is based on a semi-parametric likelihood, which is able to deal with censored data and assumes that the hazard function *h*(*t*) at time *t* is:

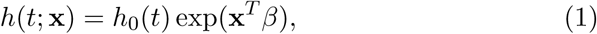

where *β* = (*β*_1_, *…, β*_*p*_) are the unknown regression coefficients, which represent the covariate effect in the survival, *h*_0_(*t*) represents the baseline hazards and **x** = (*x*_1_, *…, x*_*p*_) is the covariate vector associated to an individual.

The Cox regression model is called a semi-parametric regression model, because the baseline hazard function, *h*_0_(*t*) is not specified. This contributes for the flexibility of the model. The unknown regression coefficients, *β* are determined by maximizing the partial likelihood function

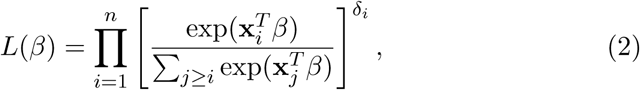

where *δ*_*i*_ is the censored indicator.

Although the Cox regression model is a widely used method due to its simplicity, the corresponding estimator has a breakdown point of 1*/n* [8], which means that the presence of outlying observations may have extreme influence on the estimation of the model parameters. In order to handle this problem, a robust version of the Cox regression model has been proposed [10] and will briefly be presented next.

### 2.2 Cox Robust regression model

The robust version of the Cox regression model [10] is based on doubly weighting the partial likelihood function of the Cox regression model.

Let *w*(*t,* **x**) be a weight function, were *w*_*ij*_ = *w*(*t*_*i*_, **x**_*j*_) and *w*_*i*_ = *w*_*ii*_ = *w*(*t*_*i*_, **x**_*i*_) are the weights for all 1 ≤ *i* ≤ *j* ≤ *n*. The solution of the unknown parameters *β* for the robust case of the partial likelihood function, presented by [10] and [11], is given by

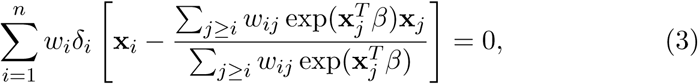

where *w*(*t,* **x**) appears in the main sum, down-weighting the uncensored observations, and in the inner sums. This allows that outlying observations will have a lower weight in the likelihood function, thus also down-weighting their influence on the parameter estimations. In this way, the most outlying observations will contribute less to the inference of the *β*.

The robust Cox is presented here as a framework that allows to infer the parameters in a more robust way when outlying observations are present, i.e. individuals that lived to long or died too early when compared to others with the same clinical conditions. Furthermore, the weights obtained with this method can give information about which observations are more influential and therefore can be considered as putative outliers.

### 2.3 Censored quantile regression

Censored quantile regression was first introduced in the econometrics literature for fixed censoring [34]. There have been many proposals in literature to overcome this issue, particularly [16] proposed the censored quantile regression as an alternative to the Cox’s regression model for survival data based on a generalization of the Kaplan-Meier one sample estimator, using recursive estimation, which relies on the assumption that the conditional quantile function is linear. An improvement of this proposal was presented in [17], where a new locally weighted censored quantile regression was considered.

The method requires that the survival time and the censoring variables are conditional independent, given the covariates, and that the quantile level of interest is linear. Due to the fact that the Cox’s regression model assumes proportional hazards, the censored quantile regression can provide more flexibility.

Let *Y*_*i*_ = min(*T*_*i*_, *C*_*i*_) represent the observed response variable, where *C*_*i*_: *i* = 1, *…, n* denoting the censoring times. The quantile regression model is defined by

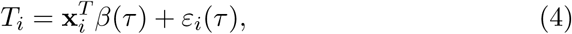

where *β*(*τ*) is a *p*-dimensional quantile coefficient vector for *τ* ∈ (0, 1), and *ε*_*i*_(*τ*) is a random error whose *τ*th conditional quantile equals zero.

Consider *Q*_*T*_*i* (*τ*|**x**_*i*_) = inf *{t*:*F* (*t*| **x**_*i*_) *≥ τ*}, the *τ* th conditional quantile of *T*_*i*_ given **x**_*i*_, and *F* (*t|***x**_*i*_) the conditional cumulative distribution function of the survival time *t* given **x**_*i*_. Then the conditional quantile is given by

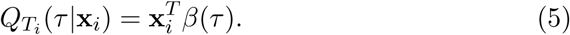

Several techniques are available in the literature to estimate the conditional quantile coefficients *β*(*τ*) (e.g. [16], [35] and [17]). As in terms of interpretation the coefficients, can be interpret as direct regression effects on the survival times. For this reason, the used of censored quantile regression must be considered when the principal interest of the study is the survival times, contractually to the Cox regression model, where the hazard rates are the main focus of the analysis.

There is a gap in the literature regarding the impact of increasing proportions of censoring on the performance of censored quantile regression. However, as referred in [5], censored quantile regression may break down with the existence of contaminated observations with extreme distance values on the predictor variables (leverage observations). For low levels of contamination, and for a central level of the quantile (*τ* = 0.5), the performance of the censored quantile regression should not be an issue.

## 3 Consensual Outlier detection in Survival Analysis

In this Section the Rank Product (RP) test is presented as a consensual technique to identify outlying observations in survival data. First several methods to detect outliers in the context of survival analysis are approached.

### 3.1 Outlier diagnostics

In the literature, three different types of unusual or discrepant cases are considered: outliers, influential and leverage observations, see [32]. An outlier can be defined as an observation with a large residual, whose dependent variable value is unusual given its value on the predictor variables; an outlier may indicate a sample peculiarity or a data entry error. A leverage observation, on the other hand, is an observation with extreme distance values on the predictor variables. An influential observation severely affects the parameters estimates, i.e., the regression coefficients change when they are removed from the dataset.

In this study, the abnormal observations that are considered are the outliers. In the context of survival analysis, an observation is considered an outlier if the individual is poorly adjusted to the model, i.e., individuals that lived too long or died too early, when compared to others with similar covariate values [2]. One of the major difference between survival data and other other types of data is the presence of censored observations. In the context of this work, it is worth stressing that the goal is not to identify the best outlier detection technique, but to delineate a method that allows to combine several approaches and generate a consensus result. In fact, given the inherent dependency of the specific methods used for outlier detection, our aim is to profit from that variability and envisage a framework that can identify aberrant observations as those consistently classified as such, independently of a particular method and/or residual definition.

Three types of outlier detection algorithms in survival analysis are described next. They are based on the residuals, the concordance c-index and, finally, on censored quantile regression.

#### 3.1.1 Residuals

Outlier diagnostics can be firstly approached via inspection of the residuals. An appropriate definition of residual is fundamental to evaluate a regression model. The usual definition of residual is the difference between the expected value of the response variable and the predicted value obtained by the model. However, when dealing with censored observations, the natural definition of residual does not stand. To overcome this difficulty, several types of residuals to analyse Cox proportional hazards models have been proposed for censored data, such as Cox-Snell, Schoenfeld, martingale, deviance, logodds and normal deviate residuals. Although, there are in the literature other types of residuals for survival data, e.g. the Schoenfeld residuals, which are very useful on the evaluation of the proportional hazards assumption, after the adjustment of a Cox model to a given dataset [13], they are not suitable for the detection of outliers.

**Cox-Snell residuals** were the first to be proposed for the proportional hazard regression model [12]. If the model is well adjusted, then the residuals should follow a known distribution.

For a given individual, **x** = (*x*_1_, *…, x*_*p*_) represents the covariate vector and *β* = (*β*_1_, *…, β*_*p*_) the regression coefficients. From the Cox regression model, the cumulative hazard function, *H*(*·*), is represented by

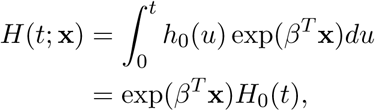

where *H*_0_(*t*) corresponds to the cumulative baseline function. The residual for the *i*^*th*^ individual, is defined as

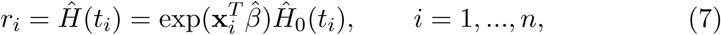

where 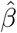 and 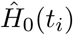 are the estimates obtained by the partial maximum likelihood of the Cox regression model. If the model is fitted correctly, the estimate values of 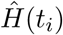 are similar to the real values of *H*(*t*_*i*_) the residuals will follow an exponential distribution with parameter *λ* = 1. That is, the residuals *r*_*i*_ should behaviour as an exponential distribution with mean value 1. Since these residuals are based on the assumption of unit exponential distribution for survival times, the expected straight line of the *H*(*·*) only applies to the exponential function [9].

Note that expression Eq. (7) does not take into account censored data. The modified Cox-Snell residuals that consider censoring are given by

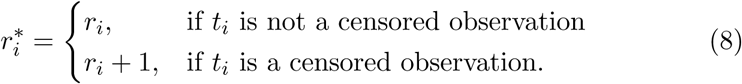

The Cox-Snell residuals are positive, and if they are higher than 1 the *t*_*i*_ observation is censored, and if the graphical representation is approximately a straight line with gradient 1 and *y*-intercept null, the model is appropriate. For more details, see [12].

**Martingale Residuals** arise from a linear transform of the Cox-Snell residuals [3] and are very useful for outlier detection. Let all the covariates be fixed, the martingale residual for the *i*^*th*^ individual is given by

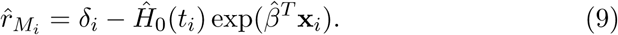

The martingale residuals are asymmetric and take values in (*-∞,* 1), which is the primary drawback of its application for outlier detection.

The martingale residuals are the difference between the observed number of the events for the *i*^*th*^ individual in (0, *t*_*i*_) and the corresponding expected number, obtained by the adjusted model. The observed number of “deaths” is one if *t*_*i*_ is not censored, i.e., is equal to *δ*_*i*_. On the other hand, *r*_*M* *i*_ is the estimate of *H*(*t*_*i*_), which can be interpret as the expected number of “deaths” in (0, *t*_*i*_), since it is only considered an individual.

The martingale residual will reveal the individuals that are not well adjusted to the model. i.e., those that lived too long (large negative values) or died too soon (values near one), when compared to other individuals with the same covariate pattern.

Although these residuals are suitable to identify outlying observations, the pronounced skewness of random departures is an inherent limitation [9]. To overcome this problem, Therneau1 and colleagues [3] introduced the deviance residuals, described next, where the asymmetry of the martingale residuals is corrected.

**Deviance Residuals** were introduced in [3], in order to overcome the fact that the martingale residuals are asymmetric distributed. The deviance residuals are defined as

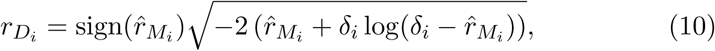

where 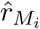 is the martingale residual for the *i*^*th*^ individual and sign(*·*) is the sign function. Notice that the deviance residuals are components of the deviance statistics, *D*, given by

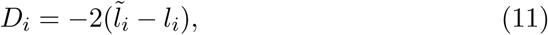

representing the difference between the log-likelihood for observation *i* under a given model 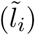 and the maximum possible log-likelihood for that observation (*l*_*i*_). In this way, 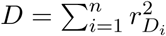, i.e., observations with high values of residuals in absolute value, are observations that are not well adjust by the model and potential outliers.

**Log-Odds and Normal deviate residuals** were proposed by [2] to identify outliers in survival analysis, measuring the differences between observed and estimated median failure time by comparing the estimated survival probability at failure time with 0.5. These residuals are quite appealing in the context of outlier detection, due to their properties and performance.

Let 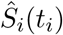 be the estimated value of the survival function for the *i*^*th*^ individual at *t*_*i*_, his observed time of failure. For a model adjusted by the Cox’s regression model, an observation is considered well predicted, if the observed survival time and its estimated median match.

The failure and non-failure of the estimated median time is coded as a binary variable following a binomial distribution. Based on this, by a transformation of the survival function using logit and probit, two residuals arise: the log-odds and normal deviate, respectively. The log-odds residual is given by

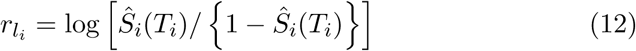

and the normal deviate residual is

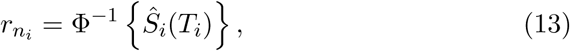

where Φ denotes the cumulative normal distribution function.

An observation *i* is considered an outlier for high absolute values of 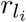 and 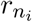. The increasing differences from a considered perfect prediction are reflected by increasing absolute values of both residuals. For more details on those residuals, see [2].

#### 3.1.2 The concordance c-index

In survival analysis, the concordance c-index [15] denotes the probability that a randomly selected subject who experienced the outcome will have a higher predictive probability of having the outcome occur compared to a randomly selected subject who did not experienced the event. Here the concordance c-index is used as a test statistics that is sensitive to the presence of outliers, i.e., the larger the number of outliers, the lower the performance of the model adjusted. Thus, the concordance c-index measures how well predicted values are concordant with the rank ordered response variables.

Three alternative methods for outlier detection in survival data based on the c-index were proposed [6,7], based on the rational that an outlier is “an observation that when absent from the data, will likely decrease the prediction error of the fitted model”.

**One-step deletion (OSD)** is an algorithm that tries to identify which observation, if removed, leads to the best improvement of the concordance c-index of the obtained model.

In the first step of the procedure, each observation of the dataset is removed temporarily, and the difference between the original c-index and the one obtained is calculated. In the following step, the most outlying case is definitively removed, and the procedure is repeated again for the remaining data.

The process ends when the quantity of erased observations reaches a pre-defined proportion of expected outliers. In the end of the algorithm the ranked list of the observations removed constitutes the putative outliers.

**Bootstrap hypothesis test (BHT)** performs *B* hypothesis tests for the concordance variation on bootstrap samples *without* the target observation *i*. Let *C*_*-i*_ be the c-index of the fitted model of the data without the *i*^*th*^ observation and *C*_0_ the c-index corresponding to the full dataset model. The hypothesis test for a certain observation *i* is:

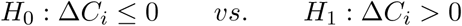

where Δ*C*_*i*_ = *C*_*-i*_ *- C*_0_. The null hypothesis states that there are no improvements on the concordance when removing observation *i*.

The algorithm starts by computing *C*_0_. For each observation *i* under test, the c-index from the model fitted without *i, C*_*-i*_, is calculated and also Δ*C*_*i*_. Then *B* bootstrap samples are generated from the data without observation *i*. The p-value is determine based on the proportion of samples having Δ*C*_*i*_ *≤* 0. The lower the p-value, the more outlying the observation is considered to be.

**Dual bootstraps hypothesis testing (DBHT)** is an improvement of the BHT described before. The BHT removes one observation from the dataset, and then evaluates the impact of each removal on the concordance c-index. Notice that the model has less observations than the original dataset, and therefore the concordance c-index has the tendency to increase, which may hamper the evaluation of the statistical significance and may increase the number of false positives.

The DBHT method generates two histograms from two opposite versions of the bootstrap procedure and compares them. In one of the samples, the observation under test is *always* removed (*A*), whereas in the other resampling scheme it is forced to be in all the (*P*) bootstrapped samples. The null hypothesis is that the expected value of 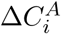 is larger than 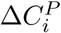 (see [6,7] for more details).

### Outlier detection in Censored quantile regression

Three different algorithms to detect outlying observations in survival data based on censored quantile regression have been proposed [4]: residual-based, boxplot and scoring. The residual-based and boxplot algorithms were developed by modifying existing ones, [2] and [18] respectively, and the scoring algorithm was introduced to provide the outlying magnitude of each point from the distribution of observations and to enable the determination of a threshold by visualizing the scores. Notice that in each run of the algorithm, the 0.5^*th*^ conditional quantile, *Q*(0.50*|***x**_*i*_), must be estimated, and its estimation is not always reliable. Consequently the results obtained by the three algorithms are not always unique. Reliable results for the estimation of *Q*(0.50*|***x**_*i*_) depends on the percentage of censoring in the dataset. Ideally the amount of censoring should not exceed 20% [38]. Next, these algorithms are briefly described.

**Residual algorithm** arises from the outlier detection algorithm for the Cox’s regression model for censored data proposed by [2] but now based on quantile regression. The residual for the *i*^*th*^ individual is given by

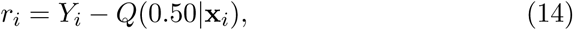

where *Q*(0.50*|***x**_*i*_) is the 50^*th*^ conditional quantile for the *i*^*th*^ individual by censored quantile regression.

Let *k*_*r*_ be a resistant parameter in order to control the cut-offs, and

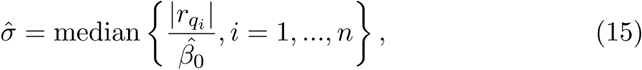

with 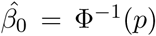 the inverse cumulative distribution of the Normal distribution for quantile *p*. Then the indicator function 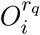 which gives the information if the *i*^*th*^ individual is or is not an outlier is defined as

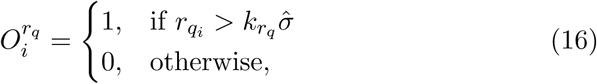

which mean that if the residual for the *i*^*th*^ individual, *r*_*q*_*i*, is higher than a threshold, 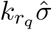 observation *i* is considered to be an outlier.

**Boxplot algorithm** is a modification of the algorithm used by [18] using quantile regression for censored data.

Obtaining the outlying individuals involves two steps. First, the censored quantile regression models are fitted for *τ* = 0.25 and *τ* = 0.75 in order to obtain the conditional quantile estimates *Q*(0.25*|***x**_*i*_) and *Q*(0.75*|***x**_*i*_), respectively. Based on those, the inter-quantile range (IQR) is determined for observation *i*, and an upper fence *UF*_*i*_ is defined as:

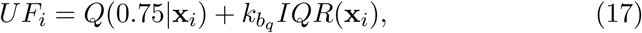

where 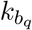 is a resistant parameter to control the tightness of cut-offs. The indicator function to declare if the *i*^*th*^ individual is an outlier is given by

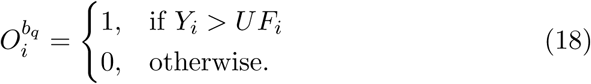

which means that an observation is considered to be an outlier if it is located above the upper fence.

**Score algorithm** In both the residual-based and the boxplot algorithms a threshold should be specified *a priori*. To overcome this limitation, the scoring algorithm was proposed, which is able to determine the deviations from the distribution of the individuals given the covariates using a flexible cut-off, 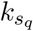.

In order to obtain the outlying individuals, first the censored quantile regression model has to be fitted for *τ* = 0.25, 0.50, 0.75 in order to obtain the conditional quantile estimates, *Q*(0.25*|***x**_*i*_), *Q*(0.50*|***x**_*i*_) and *Q*(0.75*|***x**_*i*_), respectively. By considering those, the outlying score for the *i*^th^ individual is determined by

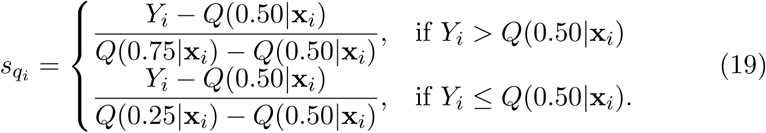

The indicator function to declare if the *i*^*th*^ by individual is an outlier is given

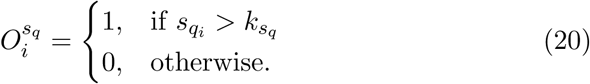

where 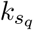 can be determined *a posteriori* by graphical visualization of the Q-Q plot of the scores.

### 3.2 Rank Product test

The outlier detection methods analysed, given their distinct assumptions and rationales, usually lead to distinct sets of solutions and outlyingness rankings. In addition, different estimated models will also significantly influence the obtained results regarding the identification of these discrepant cases.

We propose to adopt a consensus strategy to cope with this expected variability of the results, in order to delineate a more robust regression method and accurate outlier detection framework. The rationale is that, if a given observation is systematically classified as an outlier, independently of the chosen method, then our trust on the accuracy of that particular classification should increase.

Statistically, one possibility of performing a consensus ranking of the observations in terms of their relative outlyingness is to use Rank Products (RP). The required input is to have a list of all the observations ranked by their level of outlyingness, which can be based on the previously described residuals and influential measures. This non-parametric statistical technique gained great importance in detecting differential regulated genes in replicated microarray experiments [19] and can support the meta-analysis of independent studies [20]. Recently, [37] used the RP test as a consensus method to identify observations that can be influential under survival models, consequently potential outliers, in high-dimensional datasets.

Let *n* be the number of observations and *k* the number of different methods for outlier detection presented before. Consider *Z*_*ij*_ a measure of the deviance (or outlyingness) of the *i*^*th*^ observation in the *j*^*th*^ outlier detection method, with 1 ≤ *i* ≤ *n* and 1 ≤ *j* ≤ *k*. The deviance rank for each *Z*_*ij*_ considering method *j* is defined by

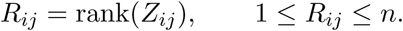

In the case of outlier detection, the lowest ranks indicate that the observation is more outlier than the others, i.e., exhibits larger deviances.

The rank product is defined by:

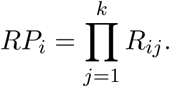

After ranked the observations by their RP, the *p*-values associated must be obtained. These *p*-values are associated with each observation under the null hypothesis that each individual ranking is uniformly distributed, which means that there each method is actually sorting randomly.

Several methods were proposed in order to estimate the statistical significance of *RP*_*i*_ under the null hypothesis of random rankings (discrete uniform distribution for each method).

In [19] the distribution of *RP*_*i*_ was based on a permutation approach. An alternative formulation that is less computational intensive was described more recently, based on an approximation of the logarithm of these values using the gamma distribution with parameters (*k,* 1) [21]. In [22] the exact probability distribution for the RP was derived. However, this approach for large *n* is increasingly expensive, which motivated another solution based on the geometric mean of upper and lower bounds, defined recursively [23]. The results shows that the algorithm provides accurate approximate *p*-values for the RP when compared to the exact ones.

Another key issue when performing these tests is related with the multiple testing problem. In fact, since many observations are tested, type-I errors (false positives) will increase. Several correction methods exist that usually adjust *α* so that the probability of observing at least one significant result due to chance remains below a desired significance level. The Bonferroni correction is one classical choice, with less conservative options also available, such as the False Discovery Rate (FDR) [24]. The FDR, which is the expected proportion of false positives among all tests that are significant, sorts in an ascendant order the p-values and divides them by their percentile rank. The measure used to determine the FDR is the q-value. For the p-value: 0.05 implies that 5% of all tests will result in false positives, instead, for the q-value: 0.05 implies that 5% of significant tests will result in false positives. In this sense the q-value is able to control the number of false discoveries in those tests. For this reason it has the ability of finding truly significant results.

The RP is used in the context of outlier detection as a consensus technique for all different results obtained by each method of outlier detection, in a model-based context.

In the next section, the RP technique is applied, where the aim is to obtain outlying observations based on some of the methods presented in Section 3.

## 4 Results and Discussion

In this section three datasets will be analysed to illustrate the performance of the outlier detection methods reviewed. A simulated dataset and two clinical case-studies, were considered. One of the issues when dealing with omics data, is the high-dimensionality problem, i.e., the number of covariates (*p*) is often much larger than the number of observations (*n*). The solution to this issue is not straightforward, with regularized optimization now standing as one of the techniques used to overcome this issue [33]. Nevertheless the aim of the present work is not the development of model variable selection methods but to evaluate ensemble outlier detection strategies. For the ovarian cancer dataset, the present proposal assumes that the variable selection in the model analysed was already performed. Regarding the myeloma dataset, where the dimensionality of the data is not an issue, the stepwise algorithm was used. The chosen datasets try to span common challenges encountered when analysing patients data, namely in survival analysis.

To use the Rank Product as a consensual technique to identify outlying observations in survival analysis, the methods considered should be independent. To guarantee this assumption, only one technique from each of the methods described in Section 3, was chosen, namely the deviance residuals (residuals), the DBHT algorithm (concordance c-index) and the Score algorithm (censored quantile regression). The deviance residual has some limitations [2], however the goal of this work is to obtain a consensual result by using the RP test, regardless of the choice of a specific technique or error definition.

To investigate the stability of the *Q*(50*|***x**_*i*_), a simulation experiment based on the real clinical data was performed. For each of the two datasets,different percentages of censoring were considered, by randomly changing the censoring indicator. In the myeloma dataset the original percentage of censored observations was 26%, and four distinct censoring values were then tested. Only for high values of censoring (> 50%) we did not obtain reliable results with the Score algorithm. Regarding the ovarian cancer dataset (45% of censored observations), the outlier detection algorithm based on censored quantile regression had good performances up to 60% censored observations. As established by [38] the amount of censoring should not exceed 20%, which is not the case for the datasets here presented (26% for the melanoma dataset and 45% for the ovarian cancer dataset). However, no convergence problems were observed when applying the algorithms, for none of the the techniques used for outlier detection. Notice also that, besides the amount censoring, also the dimension of the dataset is important to obtain reliable results. For this particular case no numerical problems were observed but it is an important aspect that should be analysied more deeply in other datasets.

All the analysis were performed in R [25]. The libraries used for were: survival, for the Cox regression model, `OutlierDC`, outlier detection method based on quantile regression, `survBootOutliers`, outlier detection based on the concordance c-index and qvalue, to determine the q-values. Two versions of the robust Cox regression model were considered: the one proposed in [10] is available in R, library coxrobust, and an improvement of this method is available in [5]. Notice that the analysis for the censored quantile was not included, due to the fact that the interest of the analysis does not rely on the importance of the survival times. The algorithm implementation to obtain the p-values for the RP, based on the geometric mean, is provided by the authors [23]. For the Cox’s robust regression model [5], an exponential weight was chosen. The number of bootstraps used for DBHT was 1000. To treat the ties in the ranks, the method used was the first occurrence wins.

All the analysis were performed in R [25] and are fully documented in the supporting information, which includes the original data, R Markdown files (Rmd Files) and HTML reports, all available at http://web.ist.utl.pt/∼susanavinga/outlierRP/.

### 4.1 Simulated data

A simulation study was carried out to highlight the importance of the RP test as a consensual technique to obtain outlying observations in survival datasets. Several datasets underlying the Cox regression model were generated, with survival times and covariates similar to real situations. For each simulated dataset a pure and contaminated models, with different parameters, were considered. For the pure model, which represents the overall tendency of the observations, the values of the parameter *β* considered were always 1. For the observations that do not follow the overall trend, the values considered for the parameter *β* ***′*** were *-*1 and *-*0.2. Two different datasets were simulated with dimensions 100 and 200, having 20 covariates each case. All covariates follow a multivariate normal distribution with zero mean, and **Σ**, the covariance matrix, equals to the identity matrix, **I**. The simulation of the survival time was based on the work of [36]. Two different models were considered to simulate the survival times: one for the pure model and other for the contaminated (outlying observations). Let *k* be the number of outlying observations considered, with *k* < *n*, the generated hazard function is given by:

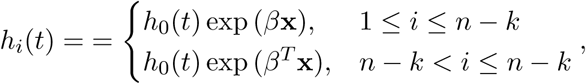

where *h*_0_(*t*) represents the baseline hazard following a Weibull distribution with scale (*λ*) and shape (*ν*) parameters. The values chosen for these parameters were 0.5, 1, 1.5. The number of outliers, *k*, was 10 for *n* = 100 and 20 for *n* = 200, representing in each case 10% of outlying observations for each simulated dataset.

The survival curves, 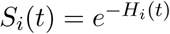, were determined based on the estimation of the *H*_*i*_(*t*), cumulative hazard function, obtained by

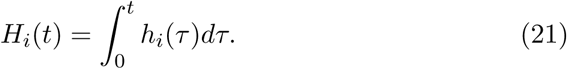

Based on the distribution obtained by generating the survival times, a censoring vector following a Bernoulli distribution, **c** = (*c*_1_, *…, c*_*n*_) was generated, with the probability of success corresponding to the proportion of censored observations. Usually the censored proportion considered is 0.2, but we also considered 0.3.

In Tables 1 and 2 are the results for the outlying observations obtained, for the simulation of 200 observations and 20% censoring. The results showed that for almost the scenarios considered, observations 181 and 186 are considered outliers regarding their *q-*value. Notice that the *q-*value (FDR) was obtained given the *p-*values ([23]) based on the RP values. As we can see for the majority of the cases, depending on the method of outlier detection chosen, different results are obtained. For instance for *β* ***′*** = *-*1,*λ* = *ν* = 1, regarding the ranks for each technique, and considering that lowest the rank more outlying observations are, observations 181 and 186 lead to different conclusions depending on each of the method chosen. However when all techniques are combined using the RP, despite the different ranks obtained for each outlier detection method, a more robust response is given. Even though the 20 outliers were not all identified (*q-*value significant), one of the advantages of using a test that combines the results of all techniques is the fact that, despite the *q-*value significant or not, we order the observations by ranks indicating their relevance for each of the methods used to detect outliers.

**Table 1:** Outlying observations for *n* = 200, 20 covariates, *β ′*= *-*1 and censoring amount of 0.2.

| | id | Rank Deviance | Rank DBHT | Rank Score | $q$-values |
| --- | --- | --- | --- | --- | --- |
| $\lambda = 1$ | 181 | 1 | 2 | 5 | 0.0010 |
| $\nu = 1$ | 186 | 3 | 1 | 11 | 0.0030 |
| $\lambda = 1.5$ | 181 | 1 | 1 | 7 | 0.0005 |
| $\nu = 0.5$ | 186 | 4 | 2 | 15 | 0.0156 |
| $\lambda = 0.5$ | 181 | 1 | 1 | 13 | 0.0021 |
| $\nu = 1.5$ | 186 | 3 | 2 | 15 | 0.0168 |

**Table 2:** Outlying observations for *n* = 200, 20 **covariates,** *β* ***′*** = *-*0.2 **and censoring amount of** 0.2.

| | id | Rank Deviance | Rank DBHT | Rank Score | $q$-values |
| --- | --- | --- | --- | --- | --- |
| $\lambda = 1$ | 186 | 3 | 1 | 80 | 0.0476 |
| $\nu = 1$ | 181 | 1 | 2 | 77 | 0.0476 |
| $\lambda = 1.5$ | 186 | 3 | 1 | 77 | 0.0424 |
| $\nu = 0.5$ | 181 | 1 | 2 | 69 | 0.0424 |
| $\lambda = 0.5$ | 183 | 1 | 1 | 124 | 0.0341 |
| $\nu = 1.5$ | 115 | 2 | 2 | 38 | 0.0341 |

### 4.2 Myeloma

The myeloma dataset is composed by clinical information on 16 covariates of 65 patients with multiple myeloma [26]. All the covariates were considered to the Cox regression model and subsequently reduced using the stepwise method in order to decrease the data dimensionality.

The results regarding the Cox regression model and the robust version are presented in Table 3. From the results, only covariate blood urea nitrogen (`bun`) is statistically significant across all the models tested, whereas gender (`sex`) and total serum protein (`total.serum.prot`) are always non significant. However, proteinuria at diagnosis (`proteinuria`) and protein in urine (`protein.urine`) are statistically significant for the Cox regression model and Cox robust proposed by [5] but not significant when using the methodology of [10]. For the hemoglobin (`hgb`), a *p*-value= 0.032 was obtained for the Cox model, but in the robust version this variable is no longer found to be significant.

**Table 3:** Results for the Cox’s regression model and Cox’s robust (both proposals) for the myeloma dataset.

| Variable | Cox |  | CoxRobust |  | CoxR (Heritier) |  |
| --- | --- | --- | --- | --- | --- | --- |
| | estimate (se) | $p$-value | estimate (se) | $p$-value | estimate (se) | $p$-value |
| hgb | -0.136 (0.063) | 0.032 | -0.138 (0.100) | 0.167 | -0.134 (0.075) | 0.075 |
| sex | -0.512 (0.363) | 0.158 | -0.546 (0.639) | 0.393 | -0.531 (0.415) | 0.201 |
| proteinuria | 0.066 (0.028) | 0.020 | 0.060 (0.055) | 0.271 | 0.059 (0.028) | 0.038 |
| protein.urine | 0.925 (0.381) | 0.015 | 0.860 (0.520) | 0.098 | 0.863 (0.404) | 0.033 |
| total.serum.prot | 0.133 (0.069) | 0.054 | 0.122 (0.063) | 0.053 | 0.123 (0.064) | 0.056 |
| bun | 0.024 (0.006) | 0.000 | 0.025 (0.007) | 0.001 | 0.025 (0.004) | 0.000 |

In order to identify outlying observations that may explain those differences, a plot of the robust estimates with log-transformed exponential weights was performed (Fig 1). Observations 40, 44 and 48 have presented the lowest weights.

**Figure 1:**
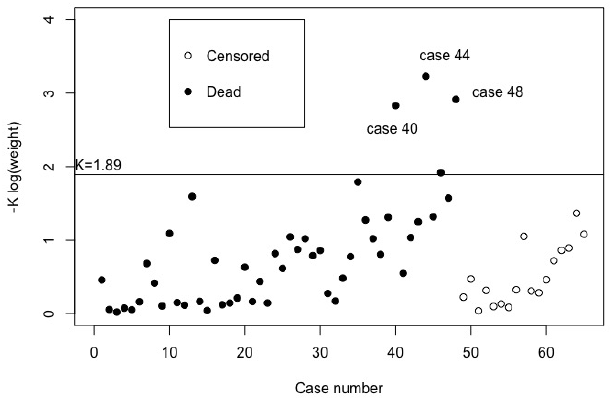
Plot of robust estimates with log-transformed exponential weight versus case number for the myeloma data with six covariates.

The results regarding the deviance residuals are presented in Fig 2. Observations 3, 2, 5 and 15 presented the highest absolute values for this residual. Different results were obtained by considered other outlier methods, such as for the concordance c-index (algorithm DBHT) and for the Score algorithm regarding the censored quantile regression.

**Figure 2:**
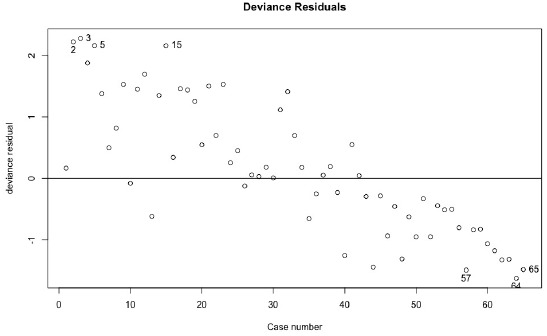
Plot of the deviance residual for the myeloma dataset with six covariates.

The top-10 outliers obtained for each method are presented in Table 4. As we can see nine of the observations are common at least for two of the methods considered: 3, 15, 21, 23, 35, 40, 44, 57 and 64. As expected, different results are obtained for each of the methods. The application of RP test allow us to combine them in a consensus ranking using the *q*-values.

**Table 4:**
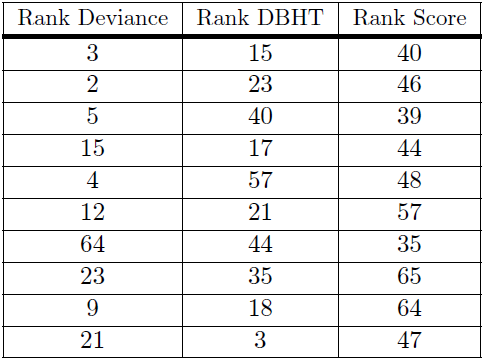
Top-10 outlying observations for the myeloma dataset with six covariates, for each technique considered.

| Rank Deviance | Rank DBHT | Rank Score |
| --- | --- | --- |
| 3 | 15 | 40 |
| 2 | 23 | 46 |
| 5 | 40 | 39 |
| 15 | 17 | 44 |
| 4 | 57 | 48 |
| 12 | 21 | 57 |
| 64 | 44 | 35 |
| 23 | 35 | 65 |
| 9 | 18 | 64 |
| 21 | 3 | 47 |

The results in Table 5 show that for the usual level of significance, none of the observations are considered outliers. Nevertheless, the proposed consensual technique is able to identify observations that should be taken under consideration for medical purpose. Notice that observation 40, which presented a lowest weight, is not considered an outlier. However it appears in the first position regarding the product of the ranks obtained by each technique. In previous studies of this dataset [26], the patient corresponding to observation 40 lived longer than the others with similar values of the covariates.

**Table 5:**
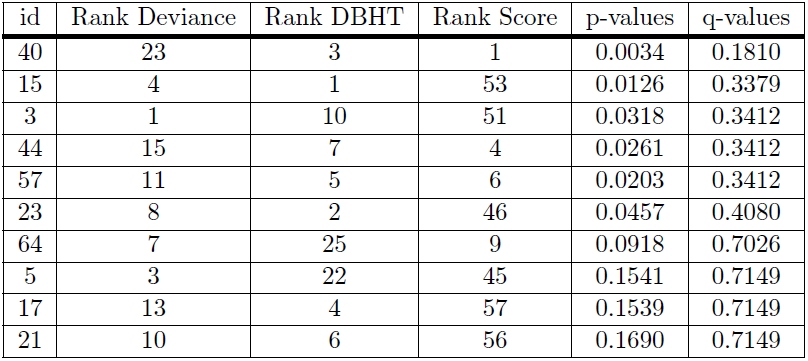
Ranks for outlier detection methods (Deviance, DBHT and Score) sorted by q-value for the myeloma dataset.

| id | Rank Deviance | Rank DBHT | Rank Score | p-values | q-values |
| --- | --- | --- | --- | --- | --- |
| 40 | 23 | 3 | 1 | 0.0034 | 0.1810 |
| 15 | 4 | 1 | 53 | 0.0126 | 0.3379 |
| 3 | 1 | 10 | 51 | 0.0318 | 0.3412 |
| 44 | 15 | 7 | 4 | 0.0261 | 0.3412 |
| 57 | 11 | 5 | 6 | 0.0203 | 0.3412 |
| 23 | 8 | 2 | 46 | 0.0457 | 0.4080 |
| 64 | 7 | 25 | 9 | 0.0918 | 0.7026 |
| 5 | 3 | 22 | 45 | 0.1541 | 0.7149 |
| 17 | 13 | 4 | 57 | 0.1539 | 0.7149 |
| 21 | 10 | 6 | 56 | 0.1690 | 0.7149 |

### 4.3 Ovarian cancer

The ovarian cancer dataset is based on gene expression data of oncological patients and is constituted by 517 observations over 12, 042 covariates. This data was obtained from The Cancer Genome Atlas (TCGA). It comprises the follow-up time, survival status and microarray gene expression of 517 patients. The microarray data was obtained using the HG-U133A platform and contains 12, 042 gene expression levels [27]. The dataset is publicly available through R package `curatedOvarianData` and it was normalized and aggregated by the TCGA consortium allowing for the analysis to be reproducible with the original dataset.

The clinical data was cleaned using `Days to last follow-up` and `Days to death` attributes to detect inconsistencies between them. Only the cases where the number of days matched were included in the analysis. The same process was performed for the attributes `Days to death` and `Vital status`, where some cases had as status `deceased`, but a missing `Days to death`.

For the analysis 18 gene expressions were considered. The variable selection was based in a previous study of [28], so no model selection was performed. The dataset is a matrix of size 517 *×* 18, and, in this case, the only genes statistically significant after fitting Cox’s and Cox’s robust models were: *CRYAB* and *SPARC* – see Table 6). Although two gene expressions were statistically significant, for a significance level of 5%, all the 18 covariates are used in the model specification and for further analysis. Despite the fact that specification of the correct model depends on the measurement of the proper covariates, in this study our concern is to demonstrate the relevance of using a consensual test (RP) to identify outlying observations and also to maintain coherence with the previous work of [28].

**Table 6:** Results for the Cox’s regression model and Cox’s robust (both proposals) for the TCGA data with 18 genes.

| variables | Cox |  |  | CoxRobust |  |  | CoxR (Heritier) |  |  |
| --- | --- | --- | --- | --- | --- | --- | --- | --- | --- |
|  | coef | se(coef) | p-value | coef | se(coef) | p-value | coef | se(coef) | p-value |
| LPL | 0.1263 | 0.0751 | 0.0924 | 0.1011 | 0.0856 | 0.2375 | 0.1011 | 0.0717 | 0.1584 |
| IGF1 | 0.0210 | 0.0600 | 0.7266 | 0.0341 | 0.0705 | 0.6289 | 0.0340 | 0.0670 | 0.6114 |
| EDNRA | 0.0224 | 0.1227 | 0.8549 | 0.0619 | 0.2119 | 0.7704 | 0.0621 | 0.1482 | 0.6752 |
| MFAP5 | 0.0165 | 0.0482 | 0.7327 | 0.0089 | 0.0622 | 0.8865 | 0.0089 | 0.0516 | 0.8630 |
| LOX | 0.1918 | 0.1251 | 0.1254 | 0.1688 | 0.1499 | 0.2604 | 0.1690 | 0.1281 | 0.1872 |
| INHBA | -0.1432 | 0.1786 | 0.4227 | -0.1556 | 0.1895 | 0.4118 | -0.1556 | 0.1841 | 0.3978 |
| THBS2 | 0.0639 | 0.0902 | 0.4787 | 0.0863 | 0.1072 | 0.4205 | 0.0862 | 0.0908 | 0.3422 |
| ADIPOQ | -0.1256 | 0.0910 | 0.1676 | -0.0727 | 0.1047 | 0.4875 | -0.0728 | 0.1001 | 0.4667 |
| NPY | 0.0552 | 0.0496 | 0.2655 | 0.0625 | 0.0710 | 0.3785 | 0.0625 | 0.0553 | 0.2590 |
| CCL11 | -0.1296 | 0.0960 | 0.1771 | -0.1578 | 0.1212 | 0.1927 | -0.1576 | 0.1013 | 0.1197 |
| VCAN | 0.0578 | 0.1009 | 0.5664 | 0.0286 | 0.1419 | 0.8404 | 0.0286 | 0.0956 | 0.7651 |
| DCN | 0.0729 | 0.0892 | 0.4133 | 0.0791 | 0.0993 | 0.4257 | 0.0791 | 0.0976 | 0.4176 |
| TIMP3 | 0.0719 | 0.0835 | 0.3891 | 0.0775 | 0.0906 | 0.3925 | 0.0775 | 0.0881 | 0.3789 |
| **CRYAB** | 0.1092 | 0.0424 | 0.0100 | 0.1179 | 0.0544 | 0.0302 | 0.1180 | 0.0437 | 0.0069 |
| CXCL12 | 0.0204 | 0.0818 | 0.8030 | 0.0129 | 0.0962 | 0.8932 | 0.0130 | 0.0879 | 0.8826 |
| **SPARC** | -0.3811 | 0.1402 | 0.0066 | -0.3978 | 0.2020 | 0.0489 | -0.3975 | 0.1332 | 0.0029 |
| CNN1 | 0.0863 | 0.1141 | 0.4493 | 0.1313 | 0.1395 | 0.3468 | 0.1313 | 0.1341 | 0.3275 |
| FBN1 | 0.1135 | 0.1690 | 0.5018 | 0.1122 | 0.2234 | 0.6154 | 0.1116 | 0.1806 | 0.5365 |

The *CRYAB* gene codes for the crystallin alpha B chain, a protein that acts as a molecular chaperone. Its function is to bind misfolded proteins and, interestingly, some defects associated to this protein and gene have already been associated with cancer, among other diseases. In particular, a recent study [29] analysed which molecular factors could affect ovarian cancer cell apoptosis and the authors found out that there was a statistical significant association between the expression of crystallin B (CRYAB) with survival. This protein has, indeed, a negative regulation of tumor necrosis, which may explain these results.

The *SPARC* gene codes for Secreted protein acidic and rich in cysteine, a protein that appears to be a regulator of cell growth, by interaction with cytokines, the extracellular matrix and also binding calcium, copper, and several others biochemical compounds. This protein is overexpressed in ovarian cancer tissues [30], playing a central role in growth, apoptosis and metastasis. It also has been identified as a candidate therapeutic target [31].

Fig 3 shows that observations 113 and 219 are identified with the lowest weights.

**Figure 3:**
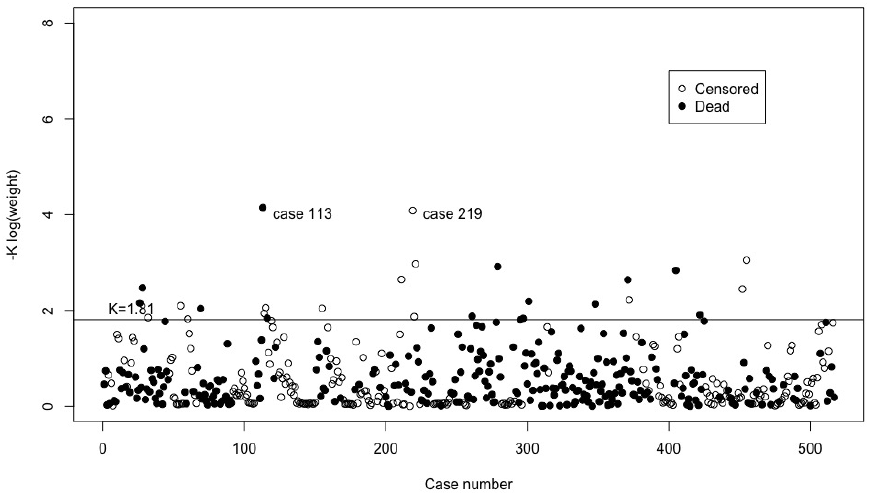
Plot of robust estimates with log-transformed exponential weight versus case number for the TCGA data with 18 genes.

Regarding the deviance residuals, the observations with the highest absolute values are 202, 346, and 415, Fig 4. The results for the top-10 most outlying observations, for the outlier detection methods chosen, are presented in Table 7, showing that observations 113, 219 and 346 are present in two of the three techniques used to identify outlying observations.

**Table 7:**
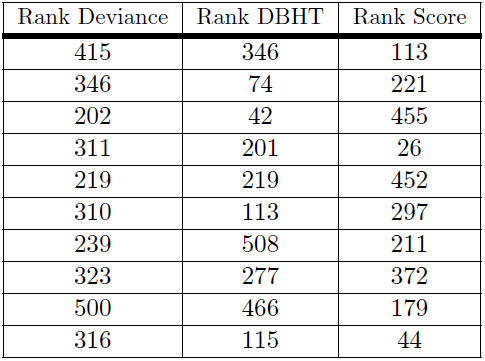
Top-10 outlying observations for the ovarian cancer with 18 genes.

**Figure 4:**
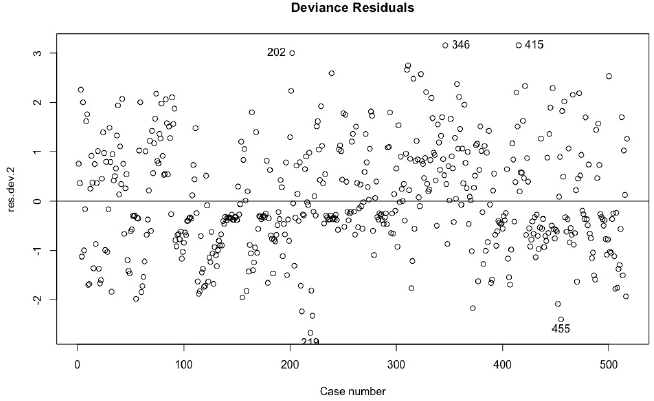
Plot of the martingale and deviance (absolute value) residuals for the TCGA data with 18 genes.

Based on the *p*-values obtained through the RP test, the observations that are considered outliers, considering the results of the *q*-values, for a 5% level of significance were: 113, 219, 221, 346 and 455 (see Table 8). Notice that observations 219 and 221 are censored representing, in the context of outliers in survival analysis, as long term survivors. On the opposite side, observation 346 represents an individual that died too early, when compared to others with similar values on the covariates.

**Table 8:** Ranks for outlier detection methods (Deviance, DBHT and Score) sorted by q-value. TCGA 18 genes dataset.

| id | Rank Deviance | Rank DBHT | Rank Score | p-values | q-values |
| --- | --- | --- | --- | --- | --- |
| 113 | 79 | 6 | 1 | 8.71E-05 | 0.0151 |
| 219 | 5 | 5 | 19 | 8.74E-05 | 0.0151 |
| 221 | 14 | 12 | 2 | 5.60E-05 | 0.0151 |
| 346 | 2 | 1 | 500 | 2.12E-04 | 0.0275 |
| 455 | 11 | 61 | 3 | 4.69E-04 | 0.0485 |
| 202 | 3 | 11 | 486 | 4.09E-03 | 0.3527 |
| 32 | 45 | 14 | 43 | 6.80E-03 | 0.3600 |
| 42 | 29 | 3 | 453 | 9.69E-03 | 0.3600 |
| 74 | 22 | 2 | 510 | 5.68E-03 | 0.3600 |
| 115 | 47 | 10 | 70 | 8.18E-03 | 0.3600 |

A cluster analysis based on the k-means algorithm was also performed in order to establish if there were any kind of relationship between those observations. By considering 4 initial clusters, observations 113 and 221, were in the same cluster. The link between cluster analysis and interpreting the results will be expanded in future work.

## 5 Conclusions

The aim of this paper was to revisit different methods for outlier detection in survival analysis and to propose a consensus method based on the RP test.

The proposed technique allows to combine the different results obtained by each method and find which observations are systematically ranked as putative outliers.

The proposed application of the RP test nevertheless illustrates that it is possible to combine disparate methods and to obtain a consensus list of putative outliers to be explored further from a clinical point of view.

By considering each method separately, none of the observations was consistently identified as an outlier for each one of the methods presented. In this sense the Rank Product test appears as a robust approach for outlier detection in the context of survival analysis.

Regarding the outlier detection method based on the concordance c-index, the fact that a resampling technique is used, different results are obtained in each run of the algorithm. Another conclusion is the fact that for a certain dataset the choice of the covariates used significantly changes the outliers identified, which may hamper a definite answer in this respect. Therefore, the results regarding the outlying observations in a given dataset are highly depended on the specific model adjusted. In this sense the consensual method here proposed is a model-based technique. It remains a question for future work how to combine this uncertainty between different Cox fittings and retrieve observations that are systematically classified as outliers independently of the model considered. Nevertheless, the RP can help to identify and filter observations that exhibit large deviances that would be expected by chance for a given number of different outlier detection methods.

Although our proposal was focused on the proportional hazards model, the method can be easily expanded to other models besides Cox regression.

In the future we attempt to use the RP to other real datasets and explore other types of regression regarding the mechanisms of outlier detection. Ensemble modelling may also provide new insights on the identification of abnormal observations in clinical data.

## Supporting information

**Rmd Files. R Markdown files with the results obtained.** The presented results are available as R Markdown (.Rmd) and html documents, along with the original data files used in the analysis performed. All these files are available at http://web.ist.utl.pt/∼susanavinga/outlierRP/.

## Acknowledgments

The authors acknowledge funding from the European Union Horizon 2020 research and innovation program under grant agreement No. 633974 (SOUND project), the Portuguese Foundation for Science & Technology (FCT), through IDMEC (under LAETA) and INESC-ID, projects UID/EMS/50022/2013, UID/CEC/50021/2013, and PERSEIDS (PTDC/EMS-SIS/0642/2014). André Veríssimo acknowledges support from FCT (SFRH/BD/97415/2013).

## References

1. Cox D.R. (1972). Regression Models and Life-Tables. Journal of the Royal Statistical Society Series B (Methodological); 34(2): 187–220.

2. Nardi A. and Schemper M. (1999). New Residuals for Cox Regression and Their Application to Outlier Screening. Biometrics; 55(2): 523–529.

3. Therneau T.M., Grambsch P.M. and Fleming T.R. (1990). Martingale-Based Residuals for Survival Models. Biometrika;77(1):147–160.

4. Eo S.H., Hong S.M. and Cho H. (2014). Identification of Outlying Observations with Quantile Regression for Censored Data. Computational Statistics; 1–17.

5. Heritier S., Cantoni E., Copt S., Victoria-Feser M.P. Robust Methods in Biostatistics. Wiley, New York. 2009.

6. Pinto J.D., Carvalho A.M. and Vinga S. (2015). Outlier Detection in Survival Analysis based on the Concordance C-index. Proceedings of the International Conference on Bioinformatics Models, Methods and Algorithms (BIOSTEC 2015); 75–82.

7. Pinto J.D., Carvalho A.M. and Vinga S. (2015). Outlier Detection in Cox Proportional Hazards Models Based on the Concordance c-Index. In: Pardalos P., Pavone M., Farinella G., Cutello V. (eds)Machine Learning, Optimization, and Big Data. Lecture Notes in Computer Science, vol 9432. Springer, Cham; 252–256. Available from: http://dx.doi.org/10.1007/978-3-319-27926-8_22.

8. Kalbeisch J.D. and Prentice R.L.. The statistical analysis of failure time data. Vol 360. John Wiley & Sons; 2011.

9. Liu X.. Survival Analysis: Models and Applications. John Wiley & Sons Ltd; United Kingdom. 2012.

10. Bednarski T. (1993). Robust Estimation in Cox’s Regression Model. Scandinavian Journal of Statistics; 20(3): 213–225.

11. Minder C.E. and Bednarski T. (1996). A robust method for proportional hazard regression. Statistics in Medicine; 15: 1033–1047.

12. Cox D.R. and Snell E.J. (1968). A General Definition of Residuals. Journal of the Royal Statistical Society Series B (Methodological); 30(2): 248–275.

13. Schoenfeld D. (1982). Partial Residuals for The Proportional Hazards Regression Model. Biometrika; 69(1): 239–2412.

14. Grambsch P.M. and Therneau T.M. (1994). Proportional Hazards tests and diagnostics based on weighted residual. Biometrika; 81: 515–526.

15. Harrell F.E., Califf R.M., Pryor D.B., Lee K.L. and Rosati R.A. (1982). Evaluating the yield of medical tests. JAMA - Journal of the American Medical Association; 247(18): 2543–2546. doi:10.1001/jama.247.18.2543.

16. Portnoy S. (2003). Censored Regression Quantiles. Journal of the American Statistical Association; 98(464): 1001–1012.

17. Wang H.J. and Wang L. (2009). Locally Weighted Censored Quantile Regression. Journal of the American Statistical Association; 104(487): 1117–1128. doi:10.1198/jasa.2009.tm08230.

18. Eo S.O., Pak D., Choi J. and Cho H. (2012). Outlier detection using projection quantile regression for mass spectrometry data with low replication. BMC Research Notes; 5(1): 236.

19. Breitling R., Armengaud P. and Herzykr P. (2004). Rank Products: A simple, yet powerful, new method to detect differentially regulated genes in replicated microarray experiments. FEBS Letters; 573: 83–92.

20. Caldas J. and Vinga S. (2014). Global Meta-Analysis of Transcriptomics Studies. PLoS One; 9(2).

21. Koziol J.A. (2010). Comments on the rank product method for analysing replicated experiments. FEBS Letters; 584: 941–944.

22. Eisinga R., Breitling R. and Heskes T. (2013). The exact probability distribution of the rank product statistics. FEBS Letters; 587: 677–682.

23. Heskes T., Eisinga R. and Breitling R. (2014). A fast algorithm for determining bounds and accurate approximate p-values of the rank product statistic for replicate experiments. BMC Bioinformatics; 15:367.

24. Storey J.D. (2002). A direct approach to false discovery rates. Journal of Royal Statistics Soc B.; 13(2): 216–225.

25. Team RC. (2012). R: A Language and Environment for Statistical Computing. Available from: http://www.R-project.org/.

26. Krall J.M., Uthoff V.A. and Hareley J.B. (1975). A step-up procedure for selecting variables associated with survival. Biometrics; 31: 49–57.

27. Network TCGAR. (2011). Integrated genomic analyses of ovarian carcinoma. Nature; 474(7353): 609–615. doi:10.1038/nature10166.

28. Zhang W., Ota T., Shridhar V., Chien J., Wu B. and Kuang R. (2013). Network-based survival analysis reveals subnetwork signatures for predicting outcomes of ovarian cancer treatment. PLoS Comput Biol.; 9(3).

29. Volkmann J., Reuning U., Rudelius M., Haefner N., Schuster T., Rose ABV, et al. (2013). High expression of crystallin B represents an independent molecular marker for unfavourable ovarian cancer patient outcome and impairs TRAIL-and cisplatin-induced apoptosis in human ovarian cancer cells. International Journal of Cancer; 132(12): 2820–2832. doi:10.1002/ijc.27975.

30. Chen J., Wang M., Xi B., Xue J., He D., Zhang J., et al. (2012). SPARC Is a Key Regulator of Proliferation, Apoptosis and Invasion in Human Ovarian Cancer. PLoS ONE; 7(8): 1–15. doi:10.1371/journal.pone.0042413.

31. Feng J., Tang L. (2014). SPARC in Tumor Pathophysiology and as a Potential Therapeutic Target. Current Pharmaceutical Design; 20(39): 6182–6190. doi:10.2174/1381612820666140619123255.

32. Nurunnabi, A.A.M., Nasser M. and Imon A.H.M.R. (2016). Identification and classification of multiple outliers, high leverage points and influential observations in linear regression. Journal of Applied Statistics; 43(3): 509–525. doi:10.1080/02664763.2015.1070806.

33. Tibshirani R. (1994). Regression Shrinkage and Selection Via the Lasso. Journal of the Royal Statistical Society, Series B; 58: 267–288.

34. Powell J.L. (1996). Censored regression quantiles. Journal of Econometrics; 32: 143–155.

35. Peng L. and Huang Y. (2008). Survival analysis with quantile regression models. Journal of the American Statistical Association; 103: 637–649.

36. Bender R., Augustin T. and Blettner M. (2005). Generating survival times to simulate Cox proportional hazard models. Statistics in Medicine; 24: 1713–1723.

37. Carrasquinha E., Verίssimo A., Lopes, M. B. and Vinga S. (2018). Identification of influential observations in high-dimensional cancer survival data through the rank product test. BioData Mining; 11(1). doi:10.1186/s13040-018-0162-z

38. Dai H. and Wang H.. Analysis for Time-to-Event Data under Censoring and Truncation, 1st edition. Academic Press. 2016.

